# Mortality Selection in a Genetic Sample and Implications for Association Studies

**DOI:** 10.1101/049635

**Authors:** Benjamin W. Domingue, Daniel W. Belsky, Amal Harrati, Dalton Conley, David Weir, Jason Boardman

## Abstract

Mortality selection is a general concern in the social and health sciences. Recently, existing health and social science cohorts have begun to collect genomic data. Causes of selection into a genomic dataset can influence results from genomic analyses. Selective non-participation, which is specific to a particular study and its participants, has received attention in the literature. But mortality selection—the very general phenomenon that genomic data collected at a particular age represents selective participation by only the subset of birth cohort members who have survived to the time of data collection—has been largely ignored. Here we test the hypothesis that such mortality selection may significantly alter estimates in polygenetic association studies of both health and non-health traits. We demonstrate mortality selection into genome-wide SNP data collection at older ages using the U.S.-based Health and Retirement Study (HRS). We then model the selection process. Finally, we test whether mortality selection alters estimates from genetic association studies. We find evidence for mortality selection. Healthier and more socioeconomically advantaged individuals are more likely to survive to be eligible to participate in the genetic sample of the HRS. Mortality selection leads to modest drift in estimating time-varying genetic effects, a drift that is enhanced when estimates are produced from data that has additional mortality selection. There is no general solution for correcting for mortality selection in a birth cohort prior to entry into a longitudinal study. We illustrate how genetic association studies using HRS data can adjust for mortality selection from study entry to time of genetic data collection by including probability weights that account for mortality selection. Mortality selection should be investigated more broadly in genetically-informed samples from other cohort studies.

## 1. Introduction

Of all individuals born in a given year, those who live until some threshold age are not a random sample. Traits, behaviors, and environmental circumstances influence survival. As cohorts age, members with disadvantageous characteristics are more likely to die off (e.g., those that smoke; Centers for Disease Control and Prevention, 2008). Such “mortality selection” can introduce bias into etiological studies of conditions that cause mortality (e.g., Vaupel & Yashin, 1985). We investigated effects of mortality selection on estimates of genetic associations in a large sample of US adults followed longitudinally during later life from the Health and Retirement Study (HRS). The HRS is among the few genetically informed databases to include longitudinal data on a population-based sample before and after the time of DNA collection. We use these longitudinal data to model mortality selection occurring between HRS baseline enrollment, which began in 1992, and DNA collection, which occurred during 2006-2008. We examined mortality selection as a function of age, socioeconomic status, health behavior, and chronic disease status.

With the information gleaned from these models of mortality, we test effects of mortality selection on two sets of genetic analyses. We tested for the influence of mortality selection on polygenic score associations with height, body mass index (BMI), educational attainment, and smoking behavior. The first set of analyses test associations between polygenic scores derived from published GWAS and the target phenotypes of those scores. The second set of analyses tests variation in those “main effects” across different HRS birth cohorts. Findings indicate that mortality selection tends to increase genetic associations with mortality-risk factors such as smoking and decrease genetic associations with protective factors such as education. With respect to changes in genetic effects across birth cohorts, mortality selection tends to increase estimates for phenotypes that have become more common in recent cohorts and decrease estimates for phenotypes that have become less common. We show that the biases introduced by mortality selection can potentially be reduced via weights from our mortality models.

### 1A. Conceptual Framework

Results from gene-environment interaction studies have made it clear that genetic effects are not constant across different environments (Hunter, 2005; Manuck & McCaffery, 2014). One potentially salient and easily measured environment is historical context as measured by birthcohort. There has been recent interest in historical context as an environmental moderator(Domingue, Conley, Fletcher, & Boardman, 2015; Guo, Liu, Wang, Shen, & Hu, 2015;Rosenquist et al., 2015), but such studies may be confounded by mortality selection. Consider the genetic effect (*G*) on some outcome (*Y*) for some birth cohort (*B*): *β*_*B*_(*G,Y*), or simply *β*_*B*_. A priori, many hypotheses about *β*_*B*_ are possible. Some genetic effects are presumably independent of birth cohort (especially over relatively small windows of time) while others are perhaps sensitive to the particular historical context during which development transpires. Whatever the behavior of *β*_*B*_, estimates of the effect, 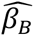, may depend upon the specific time (*T*) when data is collected. In particular, estimates of 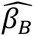 (*G,Y*) may be non-constant as a function of *T* since only those who die (indexing year of death by *D*) after the observation window (that is, *D* > *T*) are included. This is the problem of mortality selection.

The sensitivity of 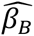 to *T* depends partially on the nature of *Y*. If mortality selection (i.e., *D* < *T*) is independent of *G* and *Y*, then missingness will not be problematic for the purposes of estimating 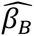 (i.e., it is MAR; Rubin & Little, 2002). For example, if development related to *Y* will largely take place after the age *T* - *B*, then this might be a reasonable hypothesis. On the other hand, if mortality selection is associated with both *G* and *Y* theninference is more challenging. Smoking, for example, is clearly associated with premature death and, as such, accurate estimates of associations from older populations may prove difficult to obtain. But any dependence will again depend upon when development occurs with respect tothe age at which respondents are studied. If development is largely complete by age *T* - *B* (e.g., has a respondent born in 1940 reported ever being a smoker by 2000 as individuals are unlikely to begin smoking after age 60), then under certain assumptions it is possible that 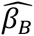 < *β*_*B*_ since observed variation in smoking behavior has been substantially reduced.

We highlight four additional issues that are relevant. First, under reasonable assumptions, 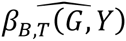 will be an unbiased estimate of *β*_*B*_(*G,Y|D* > *T*. Such estimates may be of interest in some cases (e.g., Levine & Crimmins, 2015). Bias for estimates of such effects is possible but potentially sensitive to the type of genetic effect considered. Second, the above framework could be expanded to consider age-specific genetic effects for time-varying outcomes. We do not consider such scenarios here. Third, marginal effects can be estimated as well. If variation in *β*_*B*_ as a function of birth cohort is small, these are likely to be reasonable summaries. They are also useful as first-order approximations of the genetic effect, especially when sample sizes are relatively small, and are the first types of association that we study. Finally, mortality selection is one of many selection processes that result in an individual being in a genetic sample. The consenting process, for example, also has implications for the composition of a genetic sample (e.g., McQuillan, Pan, & Porter, 2006).

### 1B. Polygenic Scores

Our examination of mortality bias in the context of genetic associations utilizes polygenic scores. Polygenic Scores (PGSs) were first introduced in 2009 (e.g., Purcell et al., 2009; Wray, Goddard, & Visscher, 2007) as flexible tools for quantifying the genetic contribution to a phenotype. Although mortality bias may is a potential concern in the study of variation at a single genetic locus, we focus on polygenic scores as they are a subject of increasing interest (e.g., Dudbridge, 2016) and are more powerful predictors of outcomes, an important consideration given the sample size available here. Also, to the extent that a PGS is a noisy measure of total additive heritability, it is an estimand of particular interest. We consider polygenic scores for smoking, educational attainment, height, and BMI that are possible due to the work of previous GWAS (Locke et al., 2015; Rietveld et al., 2013; The Tobacco and Genetics Consortium, 2010; Wood et al., 2014). These phenotypes are chosen because each has demonstrated moderate to large heritability estimates (Branigan, McCallum, & Freese, 2013; Li, Cheng, Ma, & Swan, 2003; Schousboe et al., 2003; Silventoinen et al., 2003). Further, smoking and education are each strongly related to mortality, (Hummer & Hernandez, 2013; Centers for Disease Control and Prevention, 2008).

The relationship between the remaining two phenotypes (BMI & height) and mortality is somewhat less clear. There is evidence to suggest that associations between mortality and height are mediated by other factors while the association between BMI and mortality may be more direct (Allebeck & Bergh, 1992). Other research suggests heterogeneous associations between cause-specific mortality and height (Leon, Smith, Shipley, & Strachan, 1995). Given the nature of the variants detected in the height GWAS, the lack of a clear causal role in the association between height and mortality, and the lack of a strong empirical gradient in height and early death (see our Table 2B), we hypothesize that there will be minimal bias observed for genetic associations with height. There is similar heterogeneity in cause-specific mortality and BMI (Flegal, Graubard, Williamson, & Gail, 2007) and some evidence that BMI is a poor mortality-related proxy for abdominal obesity (G. M. Price, Uauy, Breeze, Bulpitt, & Fletcher, 2006). Moreover, in our sample early mortality is associated with lower BMI. Thus, we do not expect strong bias due to mortality selection to be observed in the estimation of the genetic association with BMI. The fact that both of these phenotypes have strong genetic influences but fairly inconsistent links to mortality^1^ compared to smoking and education–which have both strong genetic underpinnings and strong links to mortality–provides an important set of comparisons as we evaluate the potential influence of mortality selection on PGS associations.

**Table 1.** Demographic characteristics of the HRS sample as a function of birth cohort and genotype status. Note: Birth cohorts are defined as: Ahead < 1924; coda 1924-1930; hrs 1931-1941; warbabies 1942-1947; early babyboomers 1948-1953; mid babyboomers 1954-1959.

|  |  |  | % Female |  | % Hispanic |  | % Black |  | % Other |  |
| --- | --- | --- | --- | --- | --- | --- | --- | --- | --- | --- |
|  | N | % Geno | Geno=Yes | Geno=No | Geno=Yes | Geno=No | Geno=Yes | Geno=No | Geno=Yes | Geno=No |
| **All** | **37319** | **33.51** | **59.07** | **54.75** | **9.07** | **12.22** | **13.12** | **20.32** | **4.76** | **7.59** |
| 0.not in any cohort | 1381 | 13.83 | 76.96 | 73.87 | 20.42 | 24.20 | 14.66 | 23.95 | 15.71 | 18.49 |
| 1.ahead | 7758 | 14.53 | 62.64 | 58.98 | 5.06 | 5.85 | 10.74 | 14.43 | 0.98 | 2.35 |
| 2.coda | 4226 | 42.81 | 55.61 | 47.91 | 5.80 | 7.36 | 8.73 | 11.42 | 3.21 | 4.14 |
| 3.hrs | 10490 | 45.72 | 55.86 | 49.79 | 8.69 | 10.20 | 13.91 | 20.39 | 3.38 | 4.74 |
| 4.warbabies | 3653 | 52.12 | 62.76 | 56.15 | 8.09 | 9.38 | 14.23 | 17.15 | 4.57 | 6.35 |
| 5.early<br>babyboomers | 4774 | 45.92 | 57.76 | 54.26 | 12.86 | 21.49 | 15.37 | 31.10 | 8.85 | 14.99 |
| 6.mid babyboomers | 5035 | 9.69 | 79.71 | 53.20 | 16.39 | 19.31 | 12.09 | 27.69 | 10.86 | 14.03 |

**Table 2.** Means of Health indicators for all HRS respondents and split by birth cohorts (panel A). Ratio of means for those lived past 2006 versus those who died before 2006 respondents in entire sample and again split by birth cohort (panel B).

| Panel A: Means | Education | BMI | Height | Smoke | CESD | Diabetes | Heart | Alzheimer's | SRH |
| --- | --- | --- | --- | --- | --- | --- | --- | --- | --- |
| **All** | **12.05** | **27.45** | **1.69** | **0.58** | **1.65** | **0.26** | **0.34** | **0.08** | **2.95** |
| 0.not in any cohort | 12.75 | 29.12 | 1.67 | 0.54 | 1.77 | 0.16 | 0.09 | 0.01 | 2.68 |
| 1.ahead | 10.71 | 24.77 | 1.67 | 0.53 | 1.87 | 0.22 | 0.52 | 0.23 | 3.32 |
| 2.coda | 11.74 | 26.49 | 1.70 | 0.61 | 1.56 | 0.29 | 0.47 | 0.13 | 3.07 |
| 3.hrs | 12.02 | 27.61 | 1.71 | 0.64 | 1.48 | 0.31 | 0.37 | 0.07 | 2.87 |
| 4.warbabies | 12.79 | 28.40 | 1.70 | 0.60 | 1.46 | 0.29 | 0.28 | 0.04 | 2.71 |
| 5.early babyboomers | 12.93 | 28.97 | 1.70 | 0.55 | 1.72 | 0.25 | 0.19 | 0.03 | 2.82 |
| 6.mid babyboomers | 12.90 | 29.50 | 1.70 | 0.57 | 1.78 | 0.20 | 0.14 | 0.01 | 2.80 |
| Panel B: Ratios | Education | BMI | Height | Smoke | CESD | Diabetes | Heart | Alzheimer's | SRH |
| **All** | **1.15** | **1.10** | **1.01** | **0.91** | **0.75** | **0.99** | **0.63** | **0.41** | **0.80** |
| 0.not in any cohort | 1.02 | 0.83 | 1.00 | 0.54 | 4.42 | Inf | Inf | 0.02 | 0.77 |
| 1.ahead | 1.07 | 1.03 | 1.00 | 0.89 | 0.81 | 1.04 | 1.05 | 1.08 | 0.88 |
| 2.coda | 1.07 | 1.02 | 0.99 | 0.77 | 0.73 | 0.83 | 0.94 | 1.25 | 0.82 |
| 3.hrs | 1.07 | 1.02 | 0.99 | 0.78 | 0.62 | 0.85 | 0.85 | 0.82 | 0.80 |
| 4.warbabies | 1.09 | 0.97 | 1.00 | 0.77 | 0.59 | 0.80 | 0.72 | 0.73 | 0.73 |
| 5.early babyboomers | 1.08 | 1.01 | 1.00 | 0.85 | 0.59 | 0.83 | 0.67 | Inf | 0.78 |
| 6.mid babyboomers | 1.03 | 1.06 | 1.00 | 0.76 | 0.95 | 0.82 | Inf | Inf | 0.86 |

## 2. Data & Methods

### 2A. Data

The Health and Retirement Study is a biennial longitudinal study starting in 1992 focused on those 50 and over. Information about work, assets, health, physical and cognitive functioning, and health care expenditures are collected. We consider 37,319 respondents in the HRS who provided any data to HRS between 1992 and 2012.^2^ We conceptualize the collection of genetic samples as a result of a two-step selection process (see inset of Figure 1). First, respondents had to live until the 2006-2008 genotyping window. In 2006, half of the sample was randomly selected to receive an enhanced interview that included saliva collection. The second half of the sample received the enhanced in-home interview in 2008. For simplicity, we focus on living until 2006 as an indicator of having avoided mortality selection.^3^ Of the original 37,319 respondents, 30,101 (80.7%) lived until at least 2006.

**Figure 1.**
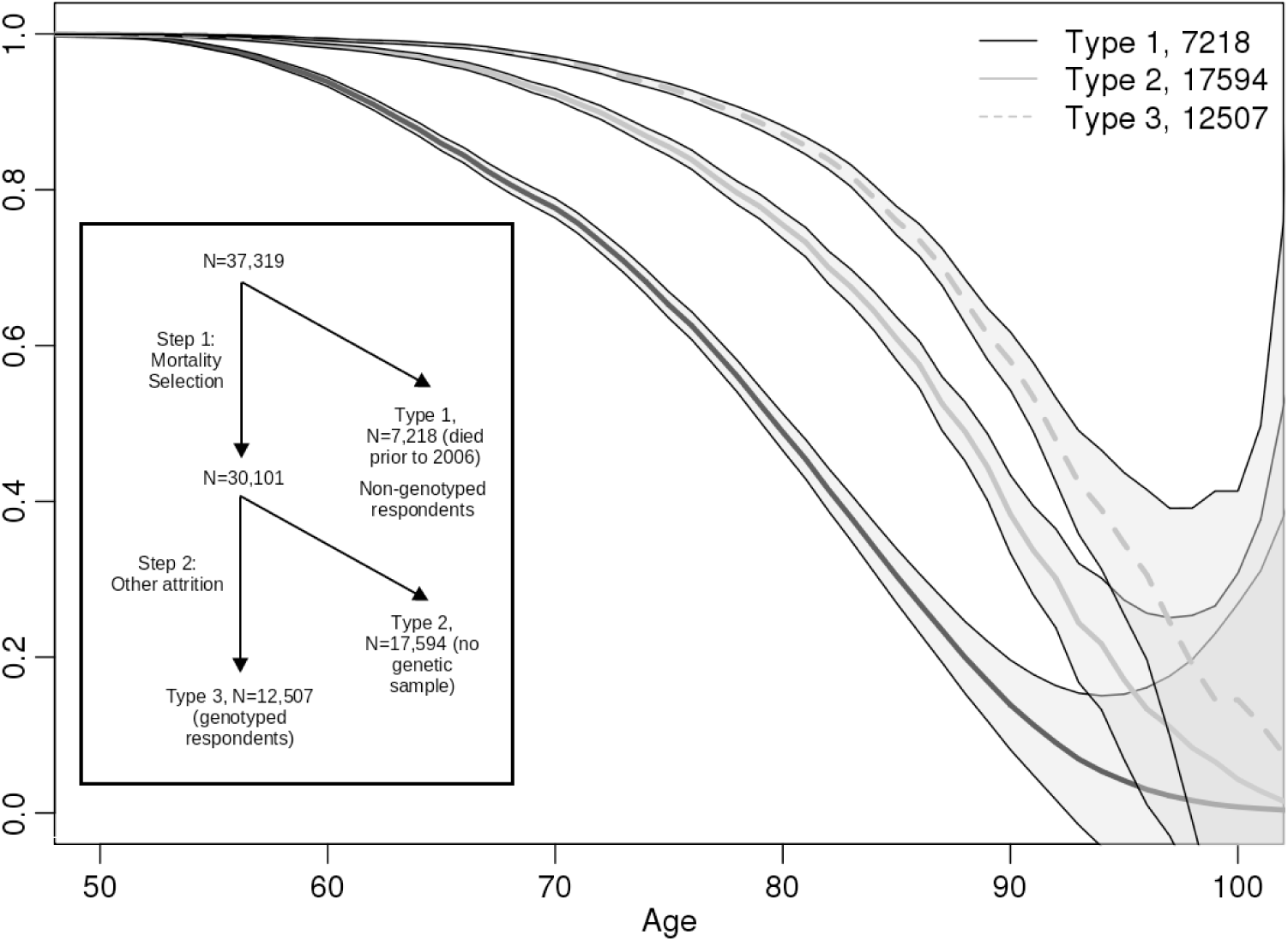
Description of two-step selection process into sample of genotyped respondents (inset). Main figure shows Kaplan-Meier survival curves for those who died prior to 2006 (type 1), those who survived through 2006 but were not genotyped (type 2) and those who were genotyped (type 3).

Second, respondents could have left the pool of candidates for genotyping due to other reasons. Respondents could be lost to HRS follow-up. Additionally, respondents who needed to be interviewed by proxy, were residing in a nursing home, or declined a face-to-face interview did not receive the enhanced interview at which the saliva collection occurred. The saliva was subsequently used for genotyping. Of the 30,101 HRS respondents who lived until 2006, 12,507 (41.6%) were genotyped in either 2006 or 2008. Although this second type of selection is not the focus of this study, we describe why genetic information is not available for 17,594 who had not died prior to 2006 in Section 2 of the SI. Figure 1 considers survival differences between those who live past 2006 but are and are not genotyped. Genotyped respondents are of course longer lived than non-genotyped respondents who lived until at least 2006, but both groups are longer lived than those who died prior to 2006. The difference between these two groups (types 2 and 3 in Figure 1) suggests that there is a secondary selection process that deserves further consideration. However, the accounting in Section 2 of the SI suggests that this is a heterogeneous group which may resist easy explanation.

Table 1 considers sex and race differences between genotyped and non-genotyped respondents in the full sample and subsamples disaggregated by birth cohort (details on birth cohort definitions are in SI Section 1). Genotyping rates were especially low for the oldest and youngest birth cohorts. In addition, while the HRS included a larger share of non-white participants in the later birth cohorts, this is not reflected in the genetically informed sample. We stratify analysis along race/sex lines to minimize the effects of these differences. We also consider a broad set of health indicators including a subjective health measure and a broader measure of socioeconomic status (years of completed education). Construction of these measures is described in the SI. Table 2 (panel A) shows the means of these variables for the entire sample and by birth cohort. Respondents had, on average, slightly more than a high school education. Roughly 58% of the sample reported ever smoking, 26% had diabetes, 34% had heart disease, 8% had Alzheimer’s or memory problems^4^, and self-reported health was roughly fair (which was coded as 3).

Genetic data for the HRS focus on single nucleotide polymorphisms (SNPs) and are based on DNA samples collected via two methods. The first phase was collected via buccal swabs in 2006 using the Quiagen Autopure method. The second phase used saliva samples collected in 2008 and extracted with Oragene. Genotype calls were then made based on a clustering of both data sets using the Illumina HumanOmni2.5-4v1 array. SNPs are removed if they are missing in more than 5% of cases, have low minor allele frequency (0.01), and are not in Hardy-Weinberg equilibrium (p<0.001). We retain approximately 1.7M SNPs after removing those that did not pass the QC filters.

### 2B. Methods

We estimate cumulative time-to-event curves by use of the Kaplan-Meier method and test via log-rank. We then use Cox proportional hazards regression analysis (Cox, 1972) to model survival and determine independent predictors,

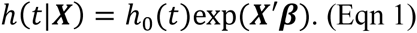

The Cox model is presented in Equation 1 and describes the association between covariates, ***X***,(genotype status, birth year, and, due to the fact that the HRS is periodically refreshed, age at first interview) and the hazard of mortalit *h(t|****X***). We also include interactions between genotype status and each of the time indicators (birth year and age at first interview). We measure duration time as the number of years between the year of the first interview and the year of the most recent interview. We also test the proportional hazards assumption; details are in Section 6 of the SI.

We model probability of inclusion in the genetic sample using logistic regression,

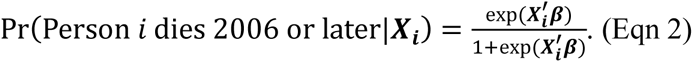

We consider various choices for the predictor matrix ***X***. Primary focus is on both a model that includes effects of birth year and the health indicators from Table 2 as well as educational attainment and a model that contains just the health indicators and education but not birth year. We also consider several alternative models to test sensitivity of the selection model to alternative assumptions. In particular, we consider models that also contain interactions between the health indicators and birth year and a model idenatified by the random forests algorithm (Liaw & Wiener, 2002), a non-parametric approach to prediction (James, Witten, Hastie, & Tibshirani, 2013).

We then use the model for mortality selection to produce probability weights (Robins,Rotnitzky, & Zhao, 1994) which we then use to create samples matched on probability of living until at least 2006 as well as in the estimation of weighted models (Lumley & others, 2004). We then utilize inverse propensity weighting to weight the observed sample to be reflective of the sample prior to mortality selection (i.e., we adjust our naïve estimate of 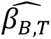, which may be biased due to mortality selection, in an attempt to more accurately recover *β*_*B*_). We stabilize probability weights (Cole & Hernán, 2008), a procedure that has been shown to yield improved estimation.

Polygenic scores were created using SNPs in the HRS genetic database that were matched to SNPs with reported results in a GWAS (we also pruned all SNPs where the risk allele identified via GWAS could not be readily identified in the HRS genetic database). For each of these SNPs, a loading was calculated as the number of trait-associated alleles multiplied by the effect-size estimated in the original GWAS. SNPs with relatively large p-values will have small effects (and thus be down weighted in creating the composite), so we do not impose a p-value threshold. Loadings were summed across the SNP set to calculate the polygenic score. The score was then standardized to have a mean of 0 and SD of 1 and then residualized across the first 10 principal components (computed amongst the non-Hispanic whites of the HRS genetic sample) to control for population stratification (A. L. Price et al., 2006). We used the second-generation PLINK software (Chang et al., 2015) for all genetic analyses.

Using the PGSs, we consider two models of genetic effects on some outcome, *Y*_*i*_. The first is a model where the genetic effect is constant across time,

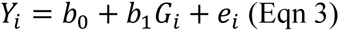

where *G*_*i*_ is individual *i*’s value for a specific PGS. The coefficient *b*_1_ from Eqn 3 is, in effect, the marginal genetic effect discussed in Section 1A. Given the nature of polygenic scores, we expect minimal bias for estimates from Eqn 3. However, we also consider models of the time-varying genetic effect, motivated by recent research on this issue (e.g., Domingue, Conley, Fletcher, & Boardman, 2015; Guo, Liu, Wang, Shen, & Hu, 2015),

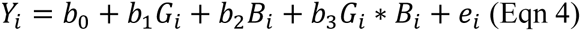

where *B*_*i*_ is the birth year of individual *i*. Eqn 4 imposes an assumption of linearity on the *β*_*B*_(*G,Y*) effect and further decomposes it into a main and interaction effect. Of primary interesthere are the estimates 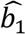 (from Eqn 3) and 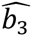 (from Eqn 4) before and after weighting by selection probability. All phenotypes are standardized within sex when used as outcomes and birth year is mean-centered. We also include sex as a control variable in the estimation of Eqns 3 and 4.

## 3. Results

### 3A. Evidence of Mortality Differences as a function of genotyping status

We first examine differences in longevity between genotyped and non-genotyped HRS respondents using non-parametric (Kaplan & Meier, 1958) survival curves, see Figure 2. In all groups, genotyped respondents live longer than non-genotyped respondents. We expand upon these results in Figures A1a and A1b of the SI finding that survival differences between the genotyped and non-genotyped samples are largest for the AHEAD, CODA, and HRS cohorts (i.e., births before 1942, which marked the start of the War Babies cohort). We next test for differences in longevity using Cox proportional hazard models (Eqn 1). Models were fit separately for men and women and for blacks and whites and included all HRS participants born 1910-1959 (N=34,669).^5^ These models show that genotyped participants are longer-lived than their non-genotyped age-peers and that this difference diminishes in later birth cohorts, see Figure 3.^6^

**Figure 2.**
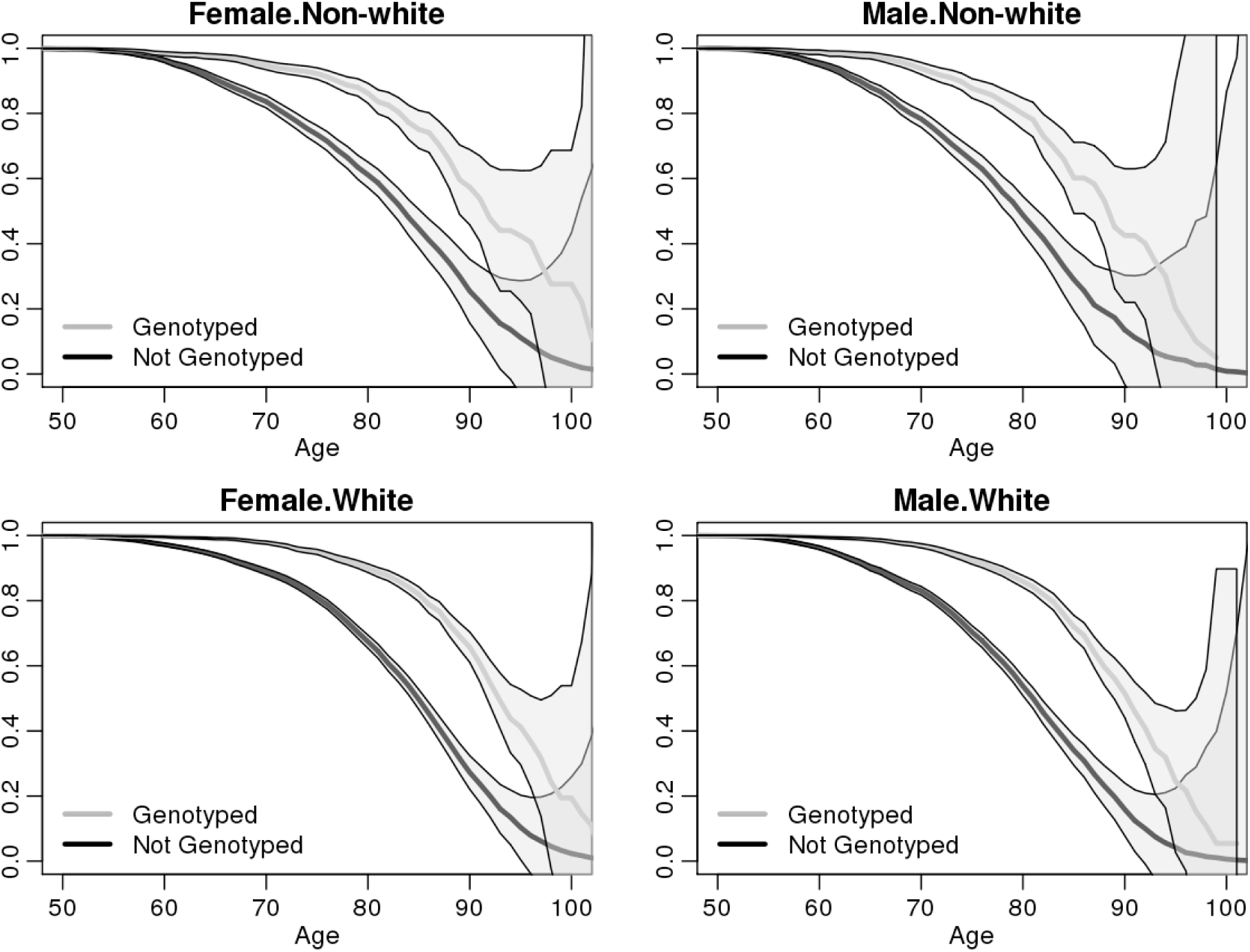
Kaplan Meier survival curves for genotyped and non-genotyped HRS respondents, split by race and sex.

**Figure 3.**
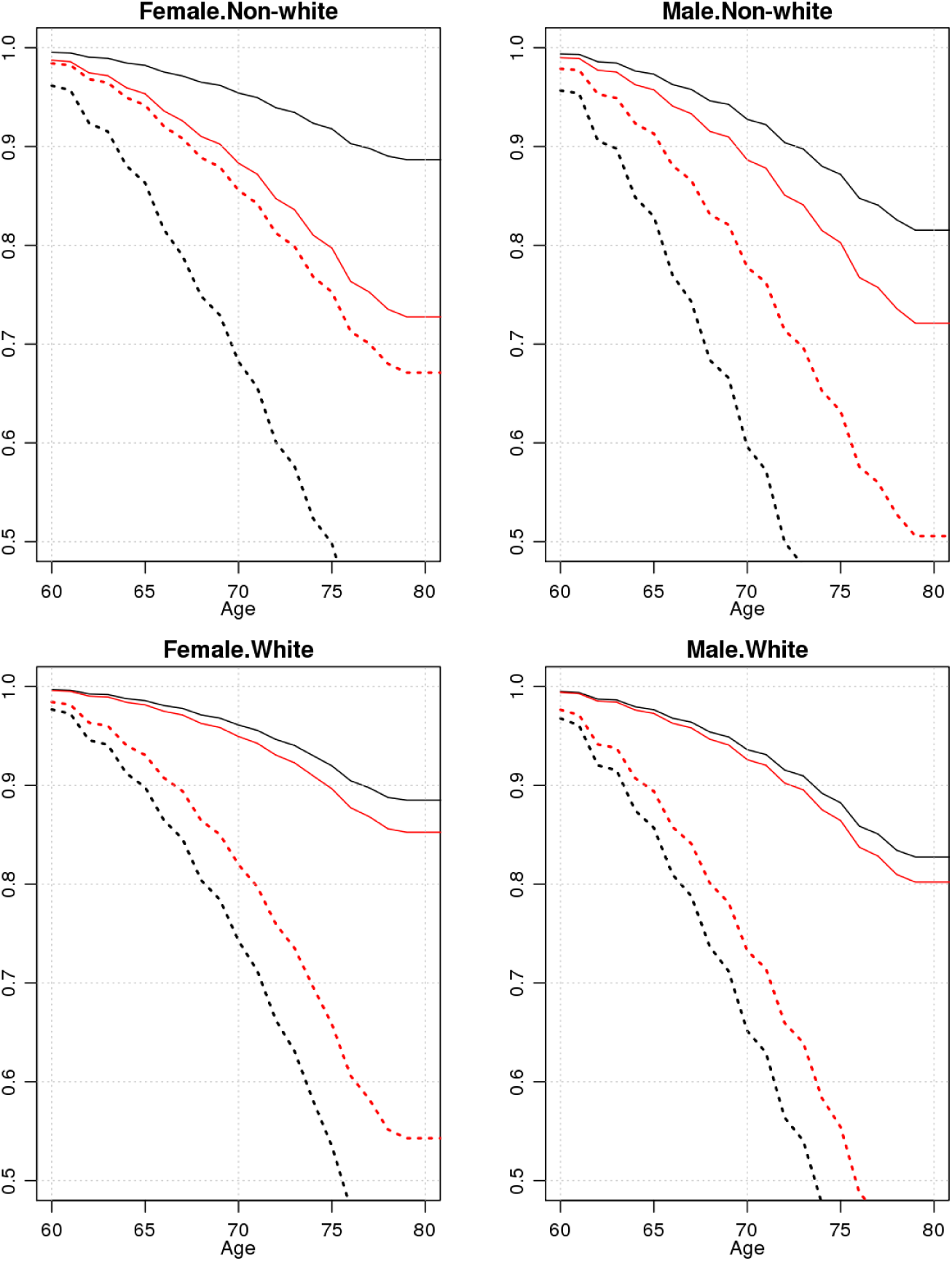
Cox survival curves for white male and female HRS respondents in the 1930 (black lines) and 1945 (red lines) birth cohorts. Solid lines show survival curves for genotyped respondents. Dashed lines show survival curves for respondents who were not genotyped. Models were estimated separately in groups defined by sex and self-reported race/ethnicity. Survival curves reflect risk for respondents aged 60 at the time of their first interview.

### 3B. Models for Mortality Selection

HRS participants who lived past 2006 were better educated and healthier as compared to non-genotyped peers (panel B of Table 2). Overall, they had completed 15% more years of education, smoked 9% less, and had dramatically lower rates of Alzheimer’s/memory problems. We used this descriptive evidence as the basis for a model of selection into the genetic sample (Eqn 2). Our analysis included all HRS participants born 1910-1959 with complete education and health data (N=30,079). Table 3 compares the fit among various models as measured by AIC and AUC. The bolded rows are our main models that include effects for all of the health conditions as well as educational attainment (of the two bolded rows, the top row includes birth year while the bottom row does not). ^7^ A consideration of the fit criteria suggests that our model including birth year does a superior job at predicting mortality. Not surprisingly, there is a substantial drop in the AUC when birth year is not included. Adding interactions between birth year and the different health conditions only improved model fit slightly (as indicated by AIC) or not at all (as indicated by AUC) and a random forest approach with the same set of predictors (including birth year) also did not yield substantial improvements in prediction of mortality.

**Table 3.** Key Estimates for models (Eqn 2) of mortality selection. Main models (bolded) include effects for all of the health conditions as well as educational attainment. Of the two bolded rows, the top row includes birth year while the bottom row does not. The “with birth year interaction” model contains interactions between all health indicators and the birth year plus the main effects from the main model. Random forests models uses a random forests algorithm based on the set of covariates used in the main model.

|  | Female Non-White |  | Male Non-White |  | Female White |  | Male White |  |
| --- | --- | --- | --- | --- | --- | --- | --- | --- |
|  | AIC | AUC | AIC | AUC | AIC | AUC | AIC | AUC |
| **Health only** | **2915** | **0.749** | **2568** | **0.728** | **7570** | **0.779** | **7090** | **0.751** |
| **Health + Birth Year** | **2248** | **0.888** | **1922** | **0.893** | **6141** | **0.879** | **5728** | **0.864** |
| W/ Birth year interaction | 2247 | 0.89 | 1922 | 0.892 | 6129 | 0.88 | 5724 | 0.864 |
| Random Forest | NA | 0.875 | NA | 0.877 | NA | 0.88 | NA | 0.862 |
| N | 5454 |  | 3967 |  | 11488 |  | 9170 |  |

### 3C. Mortality Differences in Matched Samples

We now test whether our models for selection into the genetic sample attenuate observed mortality differences between genotyped and non-genotyped respondents. We concentrate on the groups of white respondents due to sample size constraints. We utilize on an out-of-sample prediction scheme. Separately for males and females, we first select a 60% subsample of respondents and re-estimate the bolded models from Table 3. We then use the coefficient estimates to predict probability of mortality prior to 2006 in the remaining 40% sample of respondents. We then compute Cox survival models (via Eqn 1) in both the full 40% sample and a matched subsample focusing on those who were at high probability for living until at least 2006. Of interest is whether there were reduced differences in survivorship in the sampled selected for high probability of living until at least 2006. The effect of matching in this manner was dramatic, see Figure 4 (which utilizes the model including birth year). After matching for probability of early mortality (prior to 2006), the survival profiles for genotyped and non-genotyped respondents are much more comparable.

**Figure 4.**
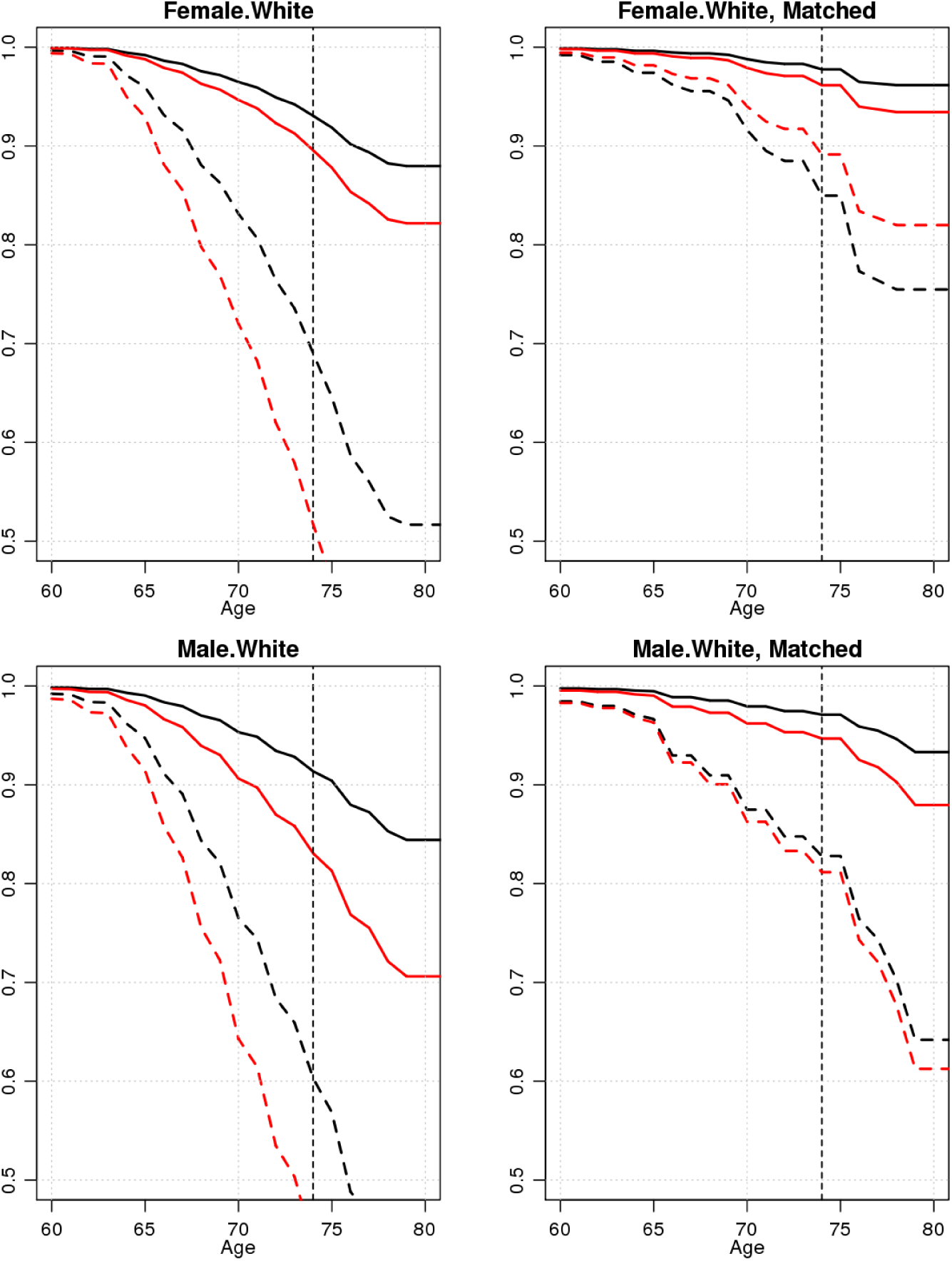
Cox survival curves for white male and female HRS respondents in the 1930 (black lines) and 1945 (red lines) birth cohorts. Solid lines show survival curves for genotyped respondents. Dashed lines show survival curves for respondents who were not genotyped. Survival curves reflect risk for respondents aged 60 at the time of their first interview. The leftside figures show survival curves based on the full sample. The right-side figures show survival curves based on a subset of genotyped and non-genotyped respondents matched according to their estimated probability of survival through 2006. The matched samples were estimated to have high survival probabilities (from the bolded model in Table 3 including birth year; estimated probabilities of genotyping were above the 70^th^ percentile as determined by the distribution of fitted probabilities amongst those who were genotyped). Since this sample was at a high probability of being genotyped, we would expect it to be healthier and do indeed see that to be the case as thicker lines are to the right of otherwise similar thinner lines.

Table 4 quantifies the increase in similarity of mortality profiles over 100 iterations of the out-of-sample prediction scheme. The table focuses on differences in survival before and after matching by considering the estimated proportion of the sample remaining after 14 years of follow-up (the time between the start of HRS and the first wave of genotyping; captured by the vertical line in Figure 4). After matching on the fitted probabilities from model including birth year, there are sizeable reductions in the survival differences between genotyped and non-genotyped respondents. Consider females born in 1930. Over all iterations, there was a raw difference in surviving proportions of 0.26 between genotyped and non-genotyped. This was reduced to a mean difference of 0.15 after matching. Reductions are even more pronounced for 1945 births for both sexes. For the model that does not include birth year, there is clearly increased similarity in mortality profiles between genotyped and non-genotyped respondents as judged by the difference in differences, although the similarity is weaker than in the model including birth year.

**Table 4.**
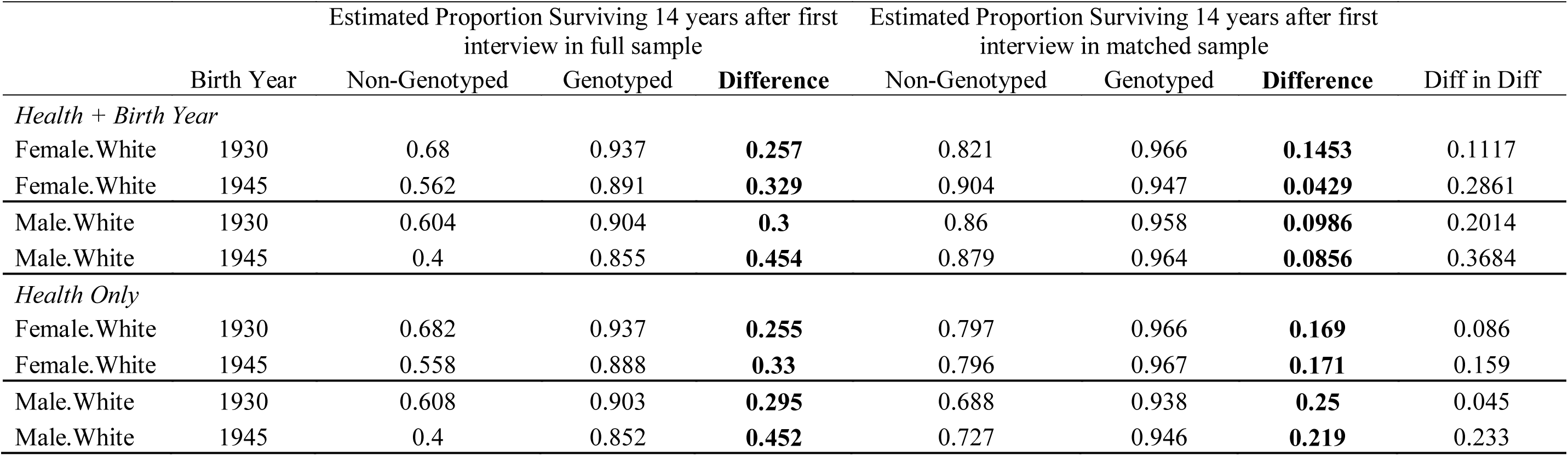
Differences in estimated survival proportions 14 years after first contact in observed and matched data (only those who had relatively high probabilities of being in the genetic sample as based on the main model). The difference in the differences was reduced by >0.15 for the 1930 births by even more for the 1945 birth cohorts. Values are based on 100 iterations of out-of-sample prediction as described in Section 3C.

### 3D. Implications for Association Studies

We first consider the mean polygenic score (focusing on BMI, height, education, and smoking) as a function of birth cohort, see Figure 5. Since the underlying distribution of the polygenic score is unlikely to have undergone substantial shifts over the relatively short time periods considered here, shifts in the means are most likely evidence for mortality selection. The most dramatic shift is for the educational attainment score. There is a substantial decline in the mean from the first observed birth cohorts to the last. Such changes may lead to bias in the resulting association estimates, a topic to which we now turn.

**Figure 5.**
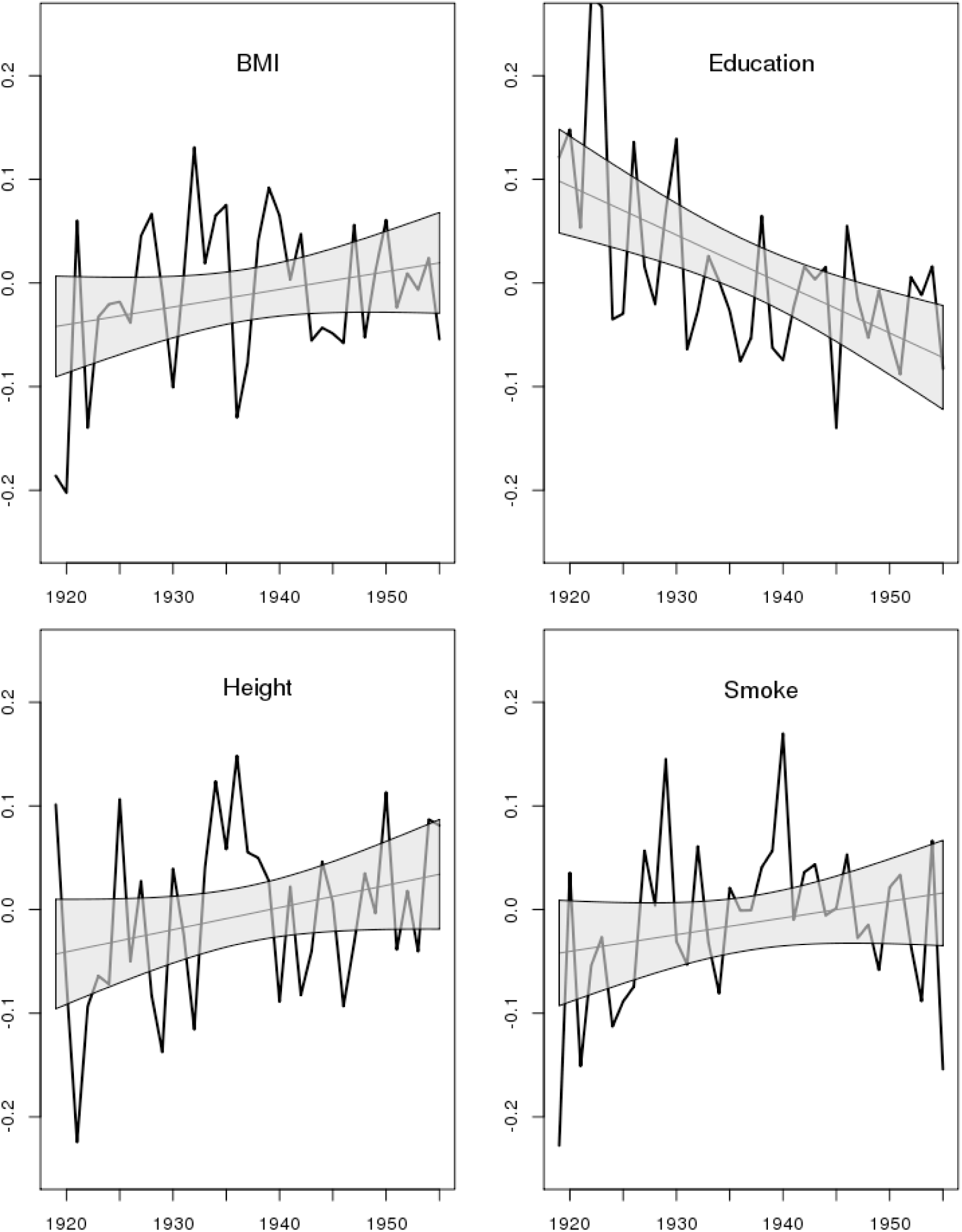
Mean polygenic score for non-Hispanic whites split by sex (N=8845) as a function of birth year (1919-1955).

We now consider the models of marginal genetic effects (Eqn 3) using polygenic scores for BMI, height, education, and smoking and their respective outcomes, see Table 5. The italicized column contains naïve (unweighted) parameter estimates using observed data. We can compare these naïve estimates to estimates from weighted models (we separately consider weights based on models that do and do not include birth year). In general, mortality selection seems to induce almost no bias in the naïve estimates as weighted estimates are fairly close to the naïve estimates (within 5% as judged by the ratios). For smoking, there is modest evidence of bias as the weighted estimate including birth year is only 95.5% of the magnitude of the original.

**Table 5.**
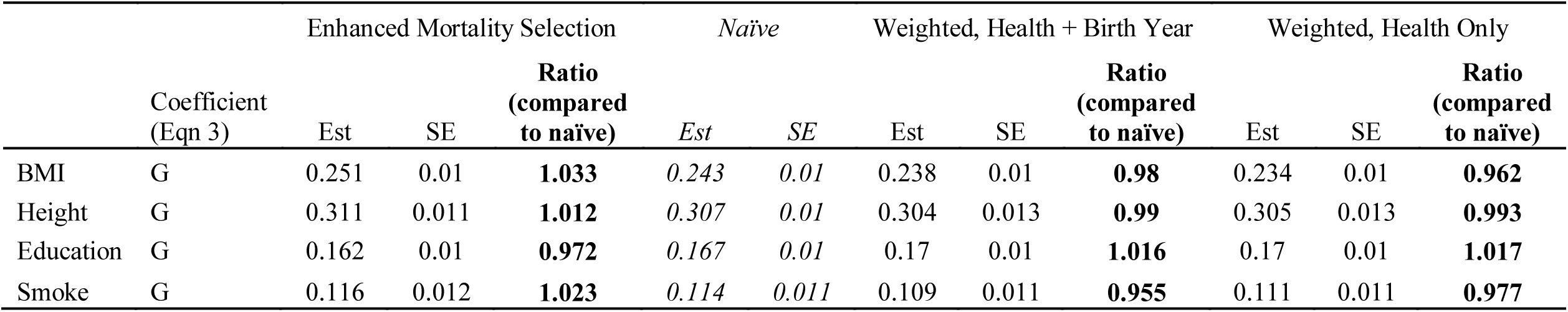
Comparison of naïve and weighted estimates of marginal associations between polygenic score and its respective trait among non-Hispanic white respondents (N=8817). The naive estimates (italicized) are unadjusted estimates of the effect of genotype on phenotype based solely on the genotyped respondents. The weighted estimates are based on identical models weighted to adjust for probability of mortality prior to 2006 (the models considered in Table 3). The ratio column is a ratio of the weighted to naïve estimate. We also consider estimates from an updated version of the HRS data that contains enhanced mortality selection (only those who lived past 2013). In addition, phenotypes are standardized within sex, main effects of gender are included but not shown, and birth year is mean-centered. All phenotypes are standardized within sex when used as outcomes.

However, we can also compare naïve and weighted estimates to estimates from a sample that contains additional mortality selection. Of the genotyped respondents, an additional 1247 respondents died after genotyping and we consider estimates based on those who have not yet died (the most recent recorded HRS deaths in our data are in 2012-2013). We emphasize that the magnitude changes in these coefficients are *opposite* the changes given by the weighting suggesting that the weighting mechanism is adjusting parameters in a sensible direction.

We then consider estimates from models of time-varying polygenic effects focusing specifically on the interaction coefficients (Table 6). Italicized columns again contain naïve parameter estimates based on observed data. For BMI and height, there is some evidence for bias in the interaction estimates. However, the potential for bias is much larger for education and smoking. For educational attainment and smoking, the naïve model seems to substantially underestimate the interaction (as compared to either set of weighted estimates). The weighted interaction estimates can also be compared to those from the data based on enhanced mortality selection. Again, for smoking and educational attainment, mortality selection leads to substantial attenuation bias in the estimated interaction coefficients.

**Table 6.**
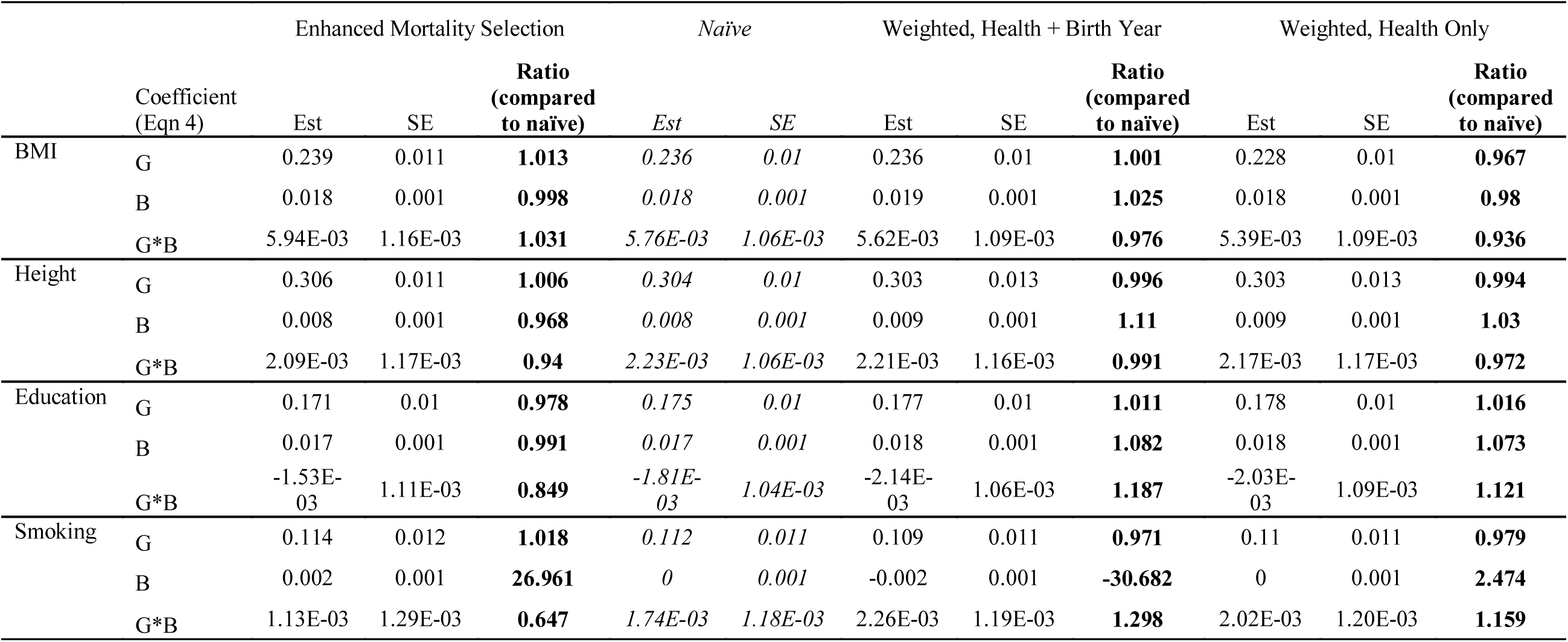
Comparison of naïve and weighted estimates of associations varying by birth cohort between a polygenic score and its respective trait among non-Hispanic white respondents (N=8817). The naive estimates (italicized) are unadjusted estimates of the effect of genotype on phenotype based solely on the genotyped respondents. The weighted estimates are based on identical models weighted to adjust for probability of mortality prior to 2006 (the models considered in Table 3). The ratio column is a ratio of the weighted to naïve estimate. We also consider estimates from an updated version of the HRS data that contains enhanced mortality selection (only those who lived past 2013). In addition, phenotypes are standardized within gender, main effects of gender are included but not shown, and birth year is mean-centered. All phenotypes are standardized within gender when used as outcomes.

## 4. Discussion

As genetic information is increasingly available in large population-based surveys, the threats to validity that traditionally apply to research based on these studies will apply to genetically informed studies as well. Here, we consider the implications of mortality selection in HRS. We demonstrated that the HRS genotyped respondents are generally healthier and more educated (see Table 2; compare to Zajacova & Burgard, 2013) as well as longer-lived (Figure 2). We then considered models for mortality selection based on the health profiles of the respondents. Samples matched on probability of mortality generally showed more similar survival profiles than unmatched samples (Figure 4), although there were still differences in the survival profiles after matching (Table 4). This suggests that generalizing association genetically informed findings from genotyped samples to larger populations may be challenging.

Our main models predicting mortality selection based on several relevant health indicators produced AUC values >0.85 (when birth year is included in the model) and >0.72 (when birth year is not included). A study on the genetic prediction of complex disease (Jostins & Barrett, 2011) suggested a maximum AUC of around 0.93 for common disease based on heritabilities, so a prediction of mortality with AUC>0.85 is a fairly accurate prediction. As a general rule, researchers should consider the association of their outcome of interest with inclusion in the genetic sample when making claims about the generalizability of their work. Depression, for example, is weakly linked to inclusion in the genetic sample via mortality selection for non-Hispanic white females. Thus, a study attempting to utilize GWAS results on depression (Ripke et al., 2013) in the HRS sample may be less problematic, in terms of generalizability, than a study utilizing GWAS results on self-reported health (Harris et al., 2015) given that the latter shows a much larger association with being in the genetic sample.

The bias observed in Tables 5/6 can be understood in terms of different mechanisms of missingness (Rubin & Little, 2002). If certain genetic profiles predispose individuals to increased risks for mortality, then individuals with such genotypes are less likely to be observed in older samples (and there is indeed evidence for this, see Figure 5). This is a form of non-ignorable missingness (MNAR) that could lead to bias in the estimated association between genotype and phenotype. Such a phenomena may explain the bias in the estimated effect of the smoking polygenic score on smoking behavior prior to weighting. However, if the genetic profile in question is largely orthogonal to mortality, then there is less reason to expect attenuation in the estimated coefficient as mortality bias is now, in effect, leading to ignorable missingness. Consider height. Results from Table 2 Panel B suggest that survivors are of the same average height as those who died prior to 2006. Thus, there is little reason to suspect bias in the estimated association between height and its polygenic score and, indeed, little is observed.

While the bias in the marginal genetic effect estimates was modest, the bias in the interaction estimates was, in some cases, quite pronounced. Moreover, we demonstrated increased bias in a sample with increased mortality selection. As interest in such interactions grows (including the gene-by-environment research that is sure to become more common as HRS releases polygenic scores), a failure to attend to this type of bias could have serious implications for many investigations. Indeed, an earlier study we conducted (Domingue et al., 2015) likely under-estimated the change in genetic influence on smoking over time due to mortality selection. Other studies (Marden et al., 2016; Rosenquist et al., 2015) considering related phenomena may also contain results that show the influence of mortality selection.

## 5. Implications for Future Research

We discuss (in order) implications for two areas of research: studies of individual variants (e.g., GWAS) and gene-environment interaction studies. The evidence presented here is focused on results from polygenic scores. However, similar concerns might be even more relevant in the study of individual genetic variants (i.e., in a GWAS) given the typical magnitudes of those effects. GWAS replication studies may consider mortality bias as a potential confounder. Again, the HRS is well-equipped to do this. For example, The HRS sample is used as a validation sample in the most recent GIANT GWAS (Locke et al., 2015) and it would be possible to ask how sensitive the replication results are to the selection concerns discussed here. The empirical evidence presented with respect to height and BMI suggest that such resulting biases may be small, but mortality bias may be more pernicious in other contexts, especially in studies of traits with strong mortality associations.

There is great interest in identifying environments which moderate genetic associations. However, mortality selection may need to be “controlled” to properly understand such associations. If selection into the genetic sample is more common among the “healthier, wealthier, and wiser” then there are theoretical reasons to expect enhanced or muted genetic associations among surviving members of the HRS. Thus, there may be influences on estimates of genetic association due to both the mechanics of mortality selection and changing structural relationships amongst the survivors and these influences may be reinforcing or opposing. This issue will merit additional attention as more genetic information continues to become available.

Finally, it is important to note that the evidence presented regarding the impact of selection on estimates of polygenic associations is limited to non-Hispanic, white adults. Our focus on non-Hispanic whites is partly due to sample size issues with estimating separate models for minority groups. But there are also important allele frequency differences across groups that may confound the observed associations with population stratification (not to mention different degrees of linkage between measured alleles and causal alleles). Furthermore, as others have noted, the lifespans of blacks and whites in the US differ by mean (roughly 3.8 years) but they also differ with respect to variation at different stages of the lifecourse (Firebaugh, Acciai, Noah, Prather, & Nau, 2014). The increased variability of lifespan in blacks may lead to less precision in our ability to fit selection probabilities for black respondents. This specific form of compression, coupled with the fact that most GWAS to date have been done with samples of European ancestry leading to weaker associations between the resulting polygenic scores and the relevant outcomes provides substantive and statistical challenges for understanding the composition of the genetic sample of non-white respondents in the HRS and how these mechanisms may be the same or different.

## Acknowledgements

This research uses data from the HRS, which is sponsored by the National Institute on Aging (Grants NIA U01AG009740, RC2AG036495, and RC4AG039029) and conducted by the University of Michigan. Research was supported by the Eunice Kennedy Shriver National Institute of Child Health and Human Development (NICHD) of the National Institutes of Health (NIH) under Award R21HD078031. Further support was provided by the NIH/NICHD-funded University of Colorado Population Center (R24HD066613). DWB is supported by an Early Career Research Fellowship from the Jacobs Foundation. AH is supported by NIH/NIA R01 AG026291.

## Footnotes

1 The relationship between mortality and BMI/height need not even be linear, although we do notconsider such a possibility here.

2 Specifically the RAND Version N data (Chien et al., 2014).

3 This simplicity comes at slight cost as there were 55 people who died in 2006 and weregenotyped, 143 in 2007, and 216 in 2008.

4 The HRS asked respondents about memory problems in waves 1-9 and replaced that questionwith one about an Alzheimer’s diagnosis in waves 10-11. We refer to “Alzheimer’s” but note theambiguity.

5 We have chosen to focus on this birth window since only 12 respondents who were born before1910 were genotyped and respondents born after 1960 would not have been categorized as beingin any of the current HRS birth cohorts.

6 The SI also considers mortality differences as a function of age at first interview, non-mortalitydifferences between genotyped and non-genotyped respondents, and survival in a restrictedsample considering only those who lived until at least 2008, see Sections 2-4.

7 Section 5 of SI contains parameter estimates from both models (with and without birth year).

